# PBF/PTTG1IP coordinates focal adhesion formation, polarity and cell motility

**DOI:** 10.64898/2026.09.14.751110

**Authors:** Merve Kocbiyik, Selvambigai Manivannan, Davina Banga, Aditi Hariharan, Mohammed E El-Asrag, Andrew D Beggs, Paloma Garcia, Dean P Larner, Saroop Raja, Jeremy A Pike, Afshan Afzal, Katie Brookes, Hannah R Nieto, Steven G Thomas, Martin L Read, Christopher J McCabe, Vicki E Smith

**Author notes:** Corresponding author (to whom reprint requests should be addressed): Dr Vicki Smith, Associate Professor in Molecular Endocrinology, University of Birmingham, UK. These authors contributed equally (Merve Kocbiyik, Selvambigai Manivannan). **Declaration of interests:** The authors declare no competing interests.

## Abstract

Directional cell migration requires cells to sense extracellular cues and coordinate adhesion, cytoskeletal remodelling and polarity, yet the molecular regulators that integrate these events remain incompletely defined. PTTG1-binding factor (PBF/PTTG1IP) is a transmembrane glycoprotein extensively characterised in pathological overexpression and cancer models, where elevated expression promotes tumourigenic cellular phenotypes. However, its endogenous physiological function remains poorly understood. Here, separate enrichment analyses of transcriptomic and phosphoproteomic profiles from PBF-overexpressing cells collectively highlighted cell adhesion, extracellular matrix organisation, cytoskeletal regulation and cell motility as major PBF-associated programmes. Using a novel Pbf knockout mouse model, we show that primary Pbf-KO mouse embryonic fibroblasts exhibit impaired migration and invasion. PBF loss reduced fibronectin adhesion, disrupted focal adhesion number, size and distribution, attenuated FAK Tyr397 phosphorylation and delayed early adhesion formation. Live-cell imaging and Golgi orientation assays further revealed altered actin organisation and impaired front–rear polarity in Pbf-KO cells. Key phenotypes were conserved in CRISPR–Cas9 PBF-KO human thyroid cancer cells, while targeted PBF re-expression restored migration in Pbf-KO MEFs. We thus identify PBF as an endogenous regulator of cell adhesion, polarity and directional motility, providing a framework for understanding how pathological PBF overexpression may promote invasive behaviour in cancer.

## INTRODUCTION

Cell migration is a fundamental physiological process requiring dynamic integration of extracellular cues, front–rear polarity, adhesion formation and disassembly, and actin cytoskeletal remodelling (1). These coordinated events are tightly regulated by multiple signalling mechanisms, including integrin-dependent activation of focal adhesion proteins and Rho GTPase-mediated control of the actin cytoskeleton (2, 3). Integrin engagement with the extracellular matrix (ECM) promotes recruitment and activation of focal adhesion proteins, including focal adhesion kinase (FAK), creating signalling complexes that couple ECM adhesion to the intracellular actin cytoskeleton (2). In parallel, Rho family GTPases regulate actin remodelling to drive membrane protrusion, contractility and polarity establishment (3). Efficient directional migration therefore depends on coordinated regulation of adhesion complexes, cytoskeletal architecture and signalling networks.

Pituitary tumor-transforming gene 1 (PTTG1)-binding factor (PBF; also known as PTTG1IP) is a ubiquitously expressed transmembrane glycoprotein that has been characterised predominantly in the context of pathological overexpression (4–7). Initially identified through its interaction with the proto-oncogene PTTG1/securin (8), PBF has subsequently been studied largely in cancer-associated settings, where its upregulation across multiple tumour types correlates with aggressive clinicopathological features (4, 9–16). Consistent with a functional role in cancer progression, increased PBF expression promotes tumourigenic phenotypes in cellular and in vivo models, including transformation, proliferation, invasion, genetic instability and tumour formation (4, 6, 10, 11, 15, 17–26).

At the molecular level, PBF has been linked to diverse processes that regulate protein localisation, signalling and cell behaviour. PBF was first shown to facilitate PTTG1 nuclear translocation (8), and later studies demonstrated roles in membrane protein trafficking through its ability to bind the sodium iodide symporter (NIS) and thyroid hormone transporter MCT8, promoting their redistribution from functional plasma membrane locations to intracellular vesicular compartments (5, 27, 28). PBF also modulates p53-dependent signalling through altered p53 protein stability and DNA damage response pathways (10, 15, 18, 19), and influences cellular proliferation in a context-dependent manner (6, 21, 23–26). In addition, PBF promotes migration and invasion in multiple cancer cell models (7, 10, 11, 15, 20, 21, 23, 26). Together, these findings suggest that PBF can influence membrane-associated signalling and cell behaviour, but whether endogenous PBF contributes to the physiological machinery that controls cell movement remains unclear.

Although previous studies provide mechanistic evidence that PBF contributes to tumourigenic phenotypes, the physiological role of endogenous PBF is largely undefined. It is unclear whether phenotypes observed in cancer-associated overexpression models reflect normal functions of endogenous PBF, or instead arise as consequences of aberrant expression. Resolving this distinction is important because cancer-associated changes in adhesion, migration and invasion frequently reflect dysregulation of cellular programmes that normally coordinate cell–environment interactions.

To determine whether endogenous PBF contributes to this cellular machinery, we generated a novel Pbf knockout mouse model and derived primary mouse embryonic fibroblasts to examine the consequences of PBF loss in a physiological cellular context. We further validated key findings using CRISPR–Cas9-mediated PBF deletion in human thyroid cells. Here, we identify PBF as an endogenous regulator of coordinated cell movement. Transcriptomic and phosphoproteomic profiling of PBF-overexpressing cells revealed molecular signatures linked to adhesion, cytoskeletal organisation and cell polarity, while loss of endogenous PBF impaired cellular adhesion, focal adhesion organisation, FAK phosphorylation, directional migration and invasion. Together, these findings establish PBF as a previously unrecognised regulator of the adhesion–polarity machinery required for efficient cell motility.

## RESULTS

### PBF overexpression identifies adhesion- and motility-associated molecular signatures

To identify cellular processes responsive to altered PBF expression, we performed parallel transcriptomic and phosphoproteomic profiling in Nthy-ori 3-1 human thyroid epithelial cells with stable PBF overexpression (**Fig. 1A**). RNA-seq identified 1,310 differentially expressed genes, comprising 625 upregulated and 685 downregulated transcripts (q < 0.1, absolute fold change ≥1.2; **Fig. 1B**; **Supplementary Table 1**). Gene Ontology (GO) enrichment analysis revealed overrepresentation of terms associated with cell adhesion, ECM organisation and cellular morphogenesis (**Fig. 1C–E**; **Supplementary Table 2**).

**Figure 1.**
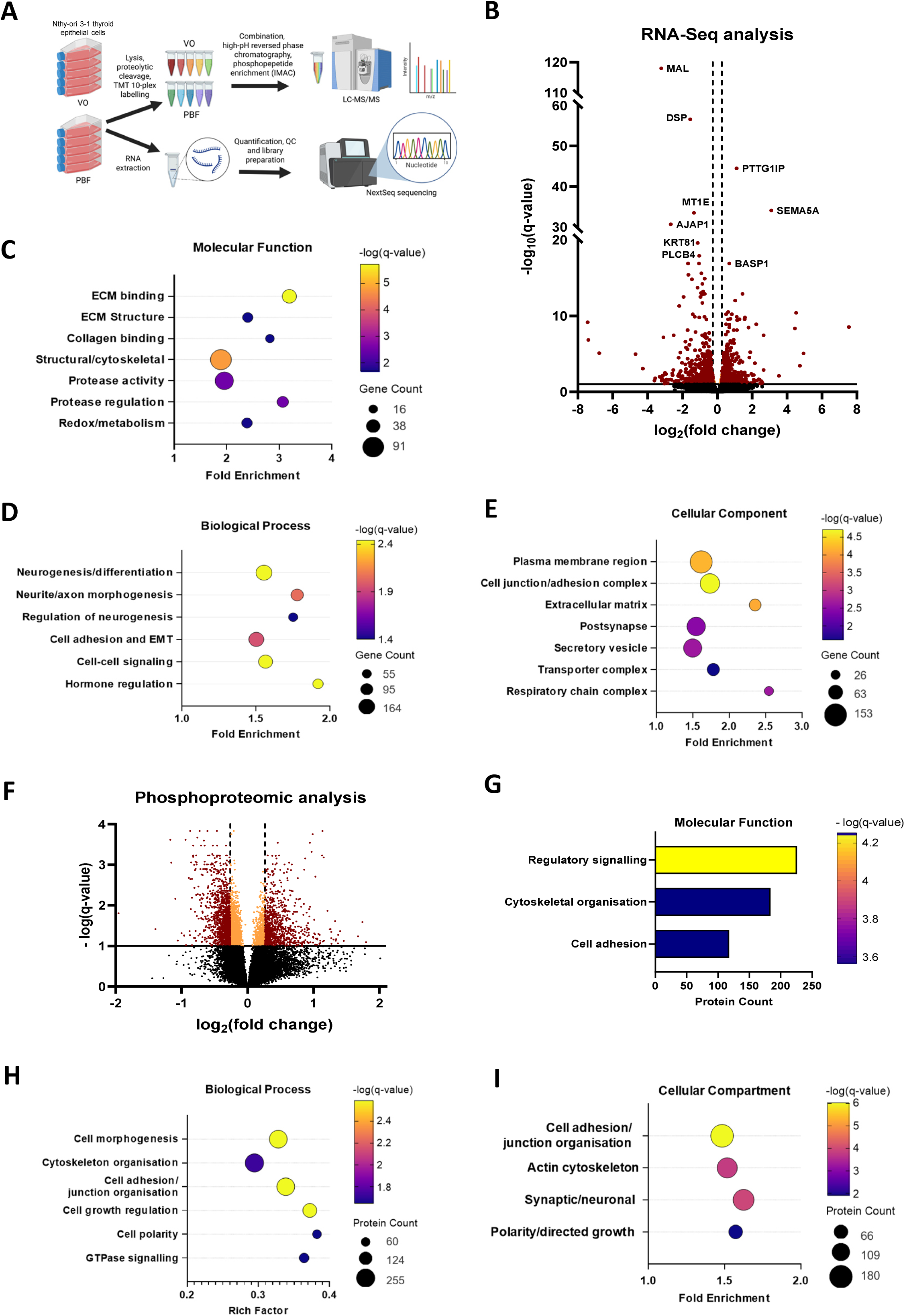
PBF overexpression identifies adhesion- and motility-associated molecular signatures. **(A)** Schematic of the experimental workflow showing stable PBF overexpression in Nthy-ori 3-1 cells followed by parallel transcriptomic and phosphoproteomic profiling. Created in BioRender. Read, M. (2026) https://BioRender.com/2p9mz2g. **(B)** Volcano plot of differentially expressed genes identified by RNA-seq following PBF overexpression. Red indicates significantly altered genes with *q* < 0.1 and absolute fold change ≥1.2. **(C–E)** Functional enrichment analysis of differentially expressed genes showing overrepresented Gene Ontology (GO) terms in the Molecular Function **(C)**, Biological Process **(D)** and Cellular Component **(E)** categories. **(F)** Volcano plot of differentially regulated phosphorylation sites following PBF overexpression. Red indicates significantly altered phosphorylation sites with *q* < 0.1 and absolute fold change ≥1.2. **(G–I)** Functional clusters of significantly overrepresented GO terms identified by enrichment analysis of differentially phosphorylated proteins in the Molecular Function **(G)**, Biological Process **(H)** and Cellular Component **(I)** categories. For all enrichment analyses, significant GO terms were defined by *q* < 0.05 following Benjamini– Hochberg false discovery rate correction. Redundant terms were consolidated into functional clusters. Circle size represents the number of unique genes or proteins within each cluster, termed the union count. Colour indicates the statistical significance of the most significantly enriched GO term within each cluster. Complete lists of significantly enriched GO terms are provided in the Supplementary Data.

Phosphoproteomic analysis identified 2,178 significantly dysregulated phosphorylation sites across 1,386 proteins, comprising 1,321 downregulated and 857 upregulated sites (q < 0.1, absolute fold change ≥1.2; **Fig. 1F**; **Supplementary Table 3**). GO enrichment analysis similarly highlighted cell adhesion, cytoskeletal organisation, cell polarity and small GTPase signalling (**Fig. 1G–I**; **Supplementary Table 4**). Together, these complementary profiling datasets highlighted related processes governing adhesion, ECM organisation, cytoskeletal regulation and polarity, providing a rationale to investigate cell motility using endogenous PBF loss-of-function models.

### Pbf loss impairs primary MEF migration and invasion

Given the enrichment of adhesion- and motility-associated signatures following PBF overexpression, together with previous evidence linking PBF to cancer cell migration and invasion (7, 10, 11, 15, 20, 21, 23, 26), we evaluated whether endogenous PBF is required for cell motility. We therefore generated a novel Pbf knockout mouse model (Pbf-KO; Pttg1ip^em1(IMPC)H^) via CRISPR–Cas9-mediated deletion of *Pbf* exon 4 (**Fig. 2A**; <u>MRC Harwell</u>). Pbf-KO mice were viable, and primary mouse embryonic fibroblasts (MEFs) were derived to assess endogenous PBF function at the cellular level.

**Figure 2.**
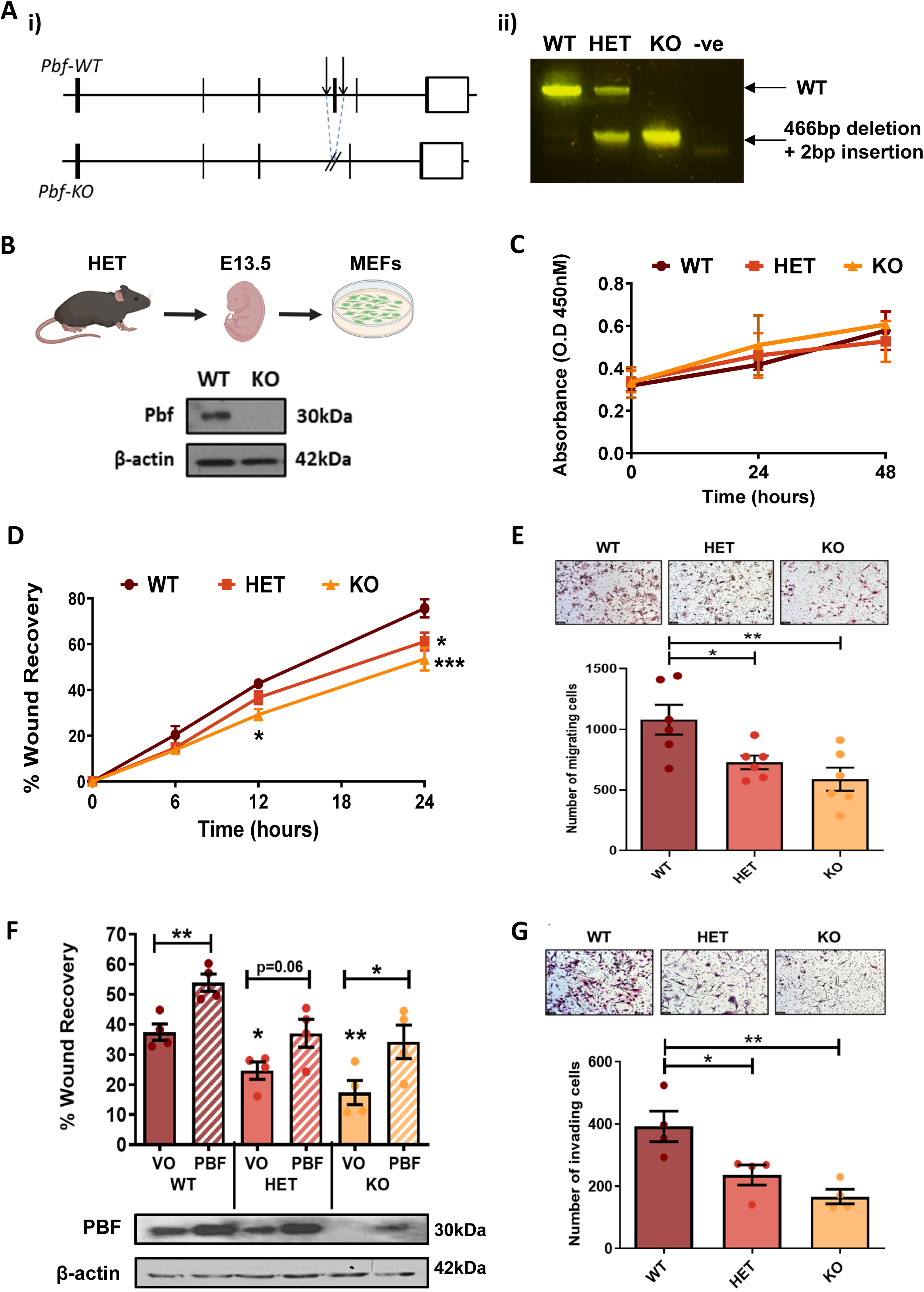
Pbf loss impairs primary MEF motility. (A) Generation and validation of the Pbf knockout mouse model. **(i)** Schematic of the murine *Pbf* gene, comprising six exons, and CRISPR–Cas9-mediated deletion of exon 4 to generate the Pbf-KO mouse. Transcript: ENSMUST00000009435.12; exon 4: ENSMUSE00001232242. **(ii)** PCR genotyping of the CRISPR–Cas9-mediated allele, comprising a 466-bp deletion and 2-bp insertion, distinguishing wild-type (WT), heterozygous (HET) and homozygous knockout (KO) MEFs. (B) Western blot confirming the loss of endogenous Pbf protein in Pbf-KO MEFs using the Cusabio anti-PBF antibody. Created in BioRender. Read, M. (2026) https://BioRender.com/r450b6v. (C) Proliferation of Pbf-WT, Pbf-HET and Pbf-KO MEFs over 48 h, assessed using the CCK-8 assay (N = 4). (D) Wound healing analysis of Pbf-WT, Pbf-HET and Pbf-KO MEFs. Wound recovery was calculated by comparing wound area at 6, 12 and 24 h with the initial wound area at 0 h (N = 4). (E) Transwell migration analysis of Pbf-WT, Pbf-HET and Pbf-KO MEFs. After 24 h, migrated cells were stained with haematoxylin and eosin and imaged at 10× magnification. Migrated cells were quantified across five fields of view per condition (N = 6). (F) Exogenous PBF expression rescues impaired migration in Pbf-deficient MEFs. Pbf-WT, Pbf-HET and Pbf-KO MEFs were transfected with either vector-only (VO) control or a PBF expression construct. A scratch wound was generated 24 h after transfection, and wound recovery was quantified after a further 24 h (N = 4). Representative Western blot showing endogenous Pbf and exogenous PBF expression in protein lysates collected 48 h after transfection, corresponding to the wound healing assay endpoint. (G) Transwell invasion analysis of Pbf-WT, Pbf-HET and Pbf-KO MEFs. After 24 h, cells that had invaded through Matrigel and traversed the membrane were stained with haematoxylin and eosin and imaged at 10× magnification. Invaded cells were quantified across five fields of view per condition (N = 4). Data are presented as mean ± SEM. Unless otherwise indicated, statistical comparisons are with Pbf-WT cells. * = *p* < 0.05, ** = *p* < 0.01, *** = *p* < 0.001.

MEFs were isolated at embryonic day 13.5 and evaluated as primary cell cultures. Loss of endogenous Pbf protein was initially confirmed by Western blotting (**Fig. 2B**). Given that PBF has been associated with a pro-proliferative phenotype in an in vivo model of thyroidal PBF overexpression (6), we first assessed cell proliferation for up to 48 h. However, no significant difference in proliferation was seen in MEFs with either heterozygous (HET) or homozygous (KO) Pbf deletion compared with wild-type (WT) MEFs (**Fig. 2C**).

Wound healing assays were used to assess the effect of Pbf deletion on cell motility. Compared with Pbf-WT MEFs, the migratory ability of Pbf-KO MEFs was significantly reduced by 31.6% and 29.3% at 12 and 24 h, respectively (**Fig. 2D**). Additionally, wound closure by Pbf-HET MEFs was significantly reduced at 24 h (**Fig. 2D**). Cell migration was also assessed using Transwell assays. Consistent with the wound healing observations, Pbf-KO MEFs exhibited a significant 45.6% reduction in cell migration compared with Pbf-WT MEFs, while Pbf-HET MEFs showed an intermediate decrease of 32.6% (**Fig. 2E**).

To determine whether impaired MEF motility was specifically attributable to Pbf loss, a rescue experiment was performed. Pbf-WT, Pbf-HET, and Pbf-KO MEFs were transfected with either vector-only (VO) control or a PBF expression construct to induce exogenous PBF expression before wound healing analysis. Western blotting confirmed marked depletion and complete loss of endogenous Pbf protein in Pbf-HET and Pbf-KO MEFs, respectively, as well as successful exogenous PBF expression (**Fig. 2F**). Initial motility data were replicated with a significant, stepwise decrease in wound recovery by 34.16% and 53.70% in Pbf-HET and Pbf-KO MEFs, respectively, compared with Pbf-WT MEFs (**Fig. 2F**). Exogenous PBF expression significantly increased wound recovery in Pbf-WT MEFs by 43.93%, indicating that increased PBF expression can enhance motility in primary cells. Importantly, exogenous PBF expression enhanced wound recovery by 33.11% and 45.19% in Pbf-HET and Pbf-KO cells, respectively, to levels that were not significantly different from Pbf-WT MEFs transfected with VO control (**Fig. 2F**).

Finally, to determine the impact of Pbf deletion on cellular invasion, Transwell invasion assays were conducted to quantify movement of Pbf-WT, Pbf-HET, and Pbf-KO MEFs through Matrigel over 24 h towards high-serum medium. A significant decrease in invasion was observed in both Pbf-KO (57.54%) and Pbf-HET (39.83%) MEFs compared with Pbf-WT cells, consistent with the migration data (**Fig. 2G**).

Taken together, these data demonstrate that Pbf loss significantly impairs the ability of primary MEFs to migrate and invade. Targeted PBF re-expression confirmed that impaired migration was specifically attributable to endogenous Pbf loss. The intermediate phenotype observed in Pbf-HET MEFs further suggests dose-dependent regulation of cell motility by PBF.

### PBF loss reduces cell adhesion

Given the impaired motility of Pbf-deficient MEFs and the enrichment of adhesion-associated terms in the PBF overexpression omics datasets, we next assessed whether PBF loss affects cell adhesion. Pbf-WT, Pbf-HET, and Pbf-KO MEFs were allowed to adhere to a fibronectin-coated surface for 1 h before removal of unattached cells. Adherent cells were fixed, stained with phalloidin (F-actin stain) and quantified. Compared with Pbf-WT MEFs, 52.7% and 43.8% fewer Pbf-HET and Pbf-KO MEFs adhered, respectively (**Fig. 3A**).

**Figure 3.**
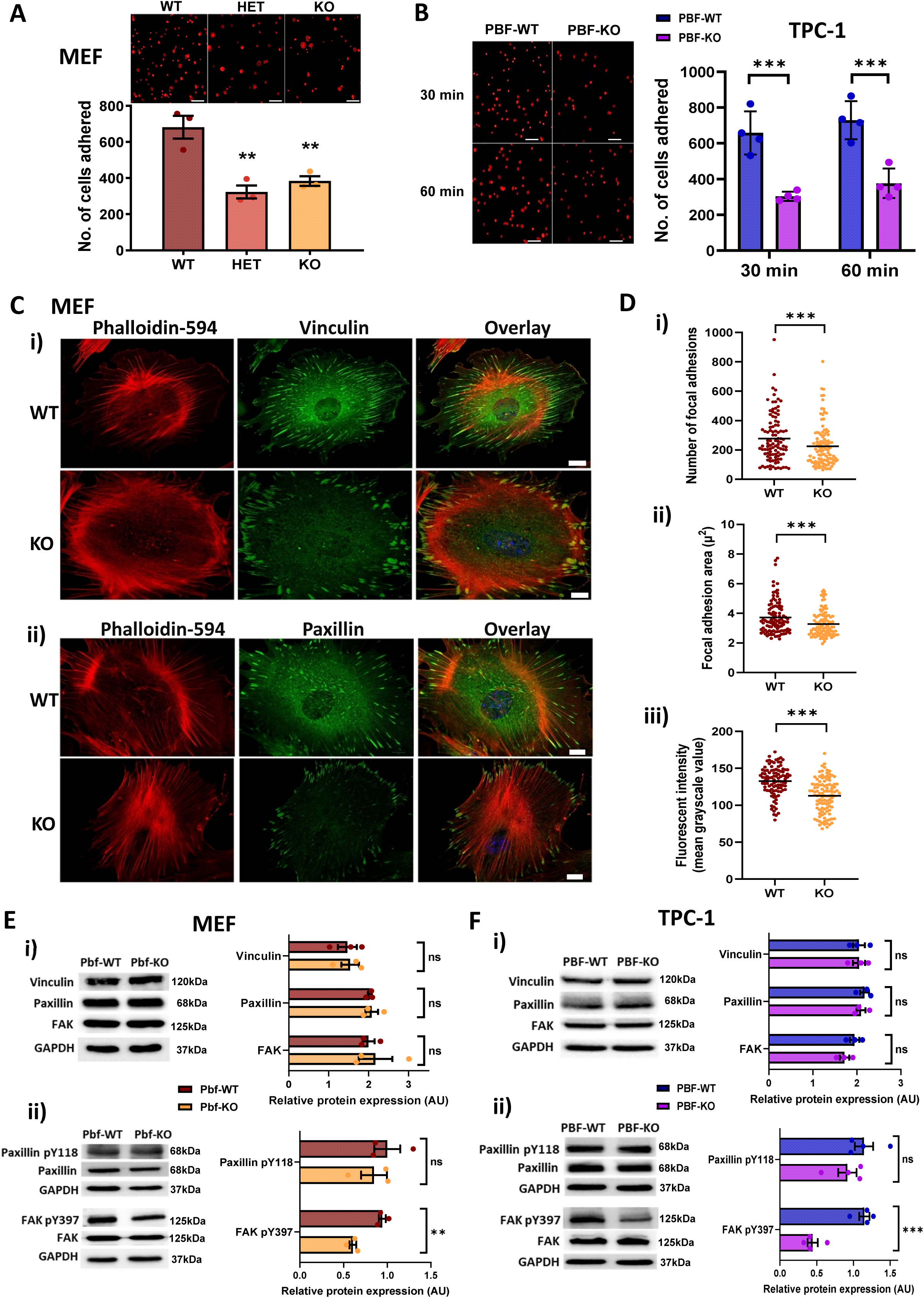
PBF loss impairs cell adhesion and disrupts focal adhesion organisation. **(A)** Adhesion of Pbf-WT, Pbf-HET and Pbf-KO MEFs to fibronectin. Cells were allowed to adhere for 1 h before removal of non-adherent cells, fixation and staining with Alexa Fluor 594–phalloidin. Representative images and quantification of adherent cells across 15 fields of view per condition are shown (N = 3). **(B)** Adhesion of parental PBF-WT and CRISPR–Cas9-mediated PBF-KO TPC-1 human papillary thyroid carcinoma cells to fibronectin after 30 min and 1 h (N = 4). **(C)** Representative immunofluorescence images of Pbf-WT and Pbf-KO MEFs stained for the focal adhesion proteins vinculin **(i)** and paxillin **(ii)** (green), together with Alexa Fluor 594–phalloidin (red) (N = 3). Scale bars, 20 μm. **(D)** Quantification of focal adhesions in vinculin-stained Pbf-WT and Pbf-KO MEFs. Focal adhesion number **(i)**, mean area in μm² **(ii)** and mean vinculin fluorescence intensity **(iii)** were quantified in 102 Pbf-WT and 108 Pbf-KO MEFs across three independent experiments. Individual cell-level measurements and overall means are shown. **(E)** Focal adhesion protein expression and phosphorylation in Pbf-WT and Pbf-KO MEFs. **(i)** Western blot analysis and quantification of total vinculin, paxillin and FAK, normalised to GAPDH (N = 3). **(ii)** Western blot analysis and quantification of paxillin-pY118 and FAK-pY397, normalised to total paxillin and FAK, respectively (N = 3). **(F)** Focal adhesion protein expression and phosphorylation in parental PBF-WT and PBF-KO TPC-1 cells. **(i)** Western blot analysis and quantification of total vinculin, paxillin and FAK, normalised to GAPDH (N = 3). **(ii)** Western blot analysis and quantification of paxillin-pY118 and FAK-pY397, normalised to total paxillin and FAK, respectively (N = 4). Data are presented as mean ± SEM. ns = not significant (*p* > 0.05), * = *p* < 0.05, ** = *p* < 0.01, *** = *p* < 0.001.

We next utilised a second cellular model of PBF deletion to assess its effect on cell adhesion, comparing the ability of CRISPR–Cas9–mediated PBF knockout TPC-1 human papillary thyroid carcinoma cells (PBF-KO) to adhere to fibronectin with that of parental TPC-1 cells (PBF-WT) (13). Significantly fewer TPC-1 PBF-KO cells adhered after 30 min (53.9%) and 60 min (48.4%) compared with PBF-WT cells (**Fig. 3B**).

These data identify PBF as an important regulator of cell–matrix adhesion in both primary MEFs and human TPC-1 thyroid cancer cells.

### PBF loss disrupts focal adhesion organisation

We next examined focal adhesion organisation in Pbf-KO MEFs by immunofluorescent staining for vinculin and paxillin (**Fig. 3C**). Both markers highlighted altered focal adhesion organisation and distribution, with adhesions appearing smaller and more radially distributed around the cell periphery in Pbf-KO MEFs compared with Pbf-WT MEFs (**Fig. 3C**). Quantification demonstrated a significant reduction in focal adhesion number, area and vinculin fluorescent staining intensity in MEFs with Pbf deletion (**Fig. 3D**).

Protein expression of key focal adhesion components was then assessed by Western blotting. Total protein levels of vinculin, paxillin and focal adhesion kinase (FAK) in Pbf-KO MEFs were not significantly different from Pbf-WT MEFs (**Fig. 3E(i)**). Tyrosine phosphorylation is an important regulatory mechanism of focal adhesion signalling and cell motility. We therefore assessed key phospho-tyrosine residues within FAK and paxillin. Following integrin engagement, FAK is activated through autophosphorylation at tyrosine 397 (Y397) (29), and can subsequently phosphorylate paxillin at tyrosine 118 (Y118) (30, 31). Phosphorylation of paxillin Y118 was comparable in Pbf-WT and Pbf-KO MEFs (**Fig. 3E(ii)**). In contrast, FAK Y397 phosphorylation was significantly reduced in Pbf-KO MEFs compared with Pbf-WT MEFs (**Fig. 3E(ii)**).

TPC-1 PBF-KO cells also demonstrated altered focal adhesion size and distribution compared with parental TPC-1 cells (**Supplementary Fig. 1**). Western blot analysis showed unaltered total protein expression of vinculin, paxillin and FAK (**Fig. 3F(i)**) and phosphorylated paxillin Y118 (**Fig. 3F(ii)**) in PBF-KO compared with PBF-WT TPC-1 cells. However, consistent with Pbf-KO MEFs, PBF-KO TPC-1 cells had significantly lower levels of phosphorylated FAK Y397 compared with parental TPC-1 cells (**Fig. 3F(ii)**).

These findings show that PBF loss reduces focal adhesion number and size, alters their cellular distribution, and attenuates phosphorylation of FAK at Y397, a key regulatory residue involved in focal adhesion signalling.

### PBF loss delays FAK phosphorylation and early focal adhesion formation

FAK Y397 phosphorylation is rapidly induced following integrin-stimulated MEF adhesion and is associated with FAK and paxillin colocalisation in early peripheral adhesions (32). Given the reduced FAK-pY397 signalling and altered focal adhesion organisation observed in Pbf-KO MEFs, we next examined whether PBF loss affects early adhesion events following attachment to fibronectin. In Pbf-WT MEFs, FAK-pY397 levels increased significantly between 15 and 30 min, indicating FAK activation (**Fig. 4A**). In contrast, Pbf-KO MEFs showed no induction of FAK-pY397, which was significantly attenuated compared with Pbf-WT MEFs at 30 min (**Fig. 4A**). In parallel, FAK and paxillin staining revealed distinct early peripheral adhesions in Pbf-WT MEFs, whereas Pbf-KO cells displayed less prominent peripheral adhesions at the same time points (**Fig. 4B**). Similarly, the significant induction of FAK-pY397 seen in PBF-WT TPC-1 cells 30 min after adhesion to fibronectin was lacking in PBF-KO cells (**Fig. 4C**), and early peripheral adhesion formation was markedly reduced at this time point (**Fig. 4D**). These data indicate that PBF loss delays early adhesion-associated FAK activation and focal adhesion formation.

**Figure 4.**
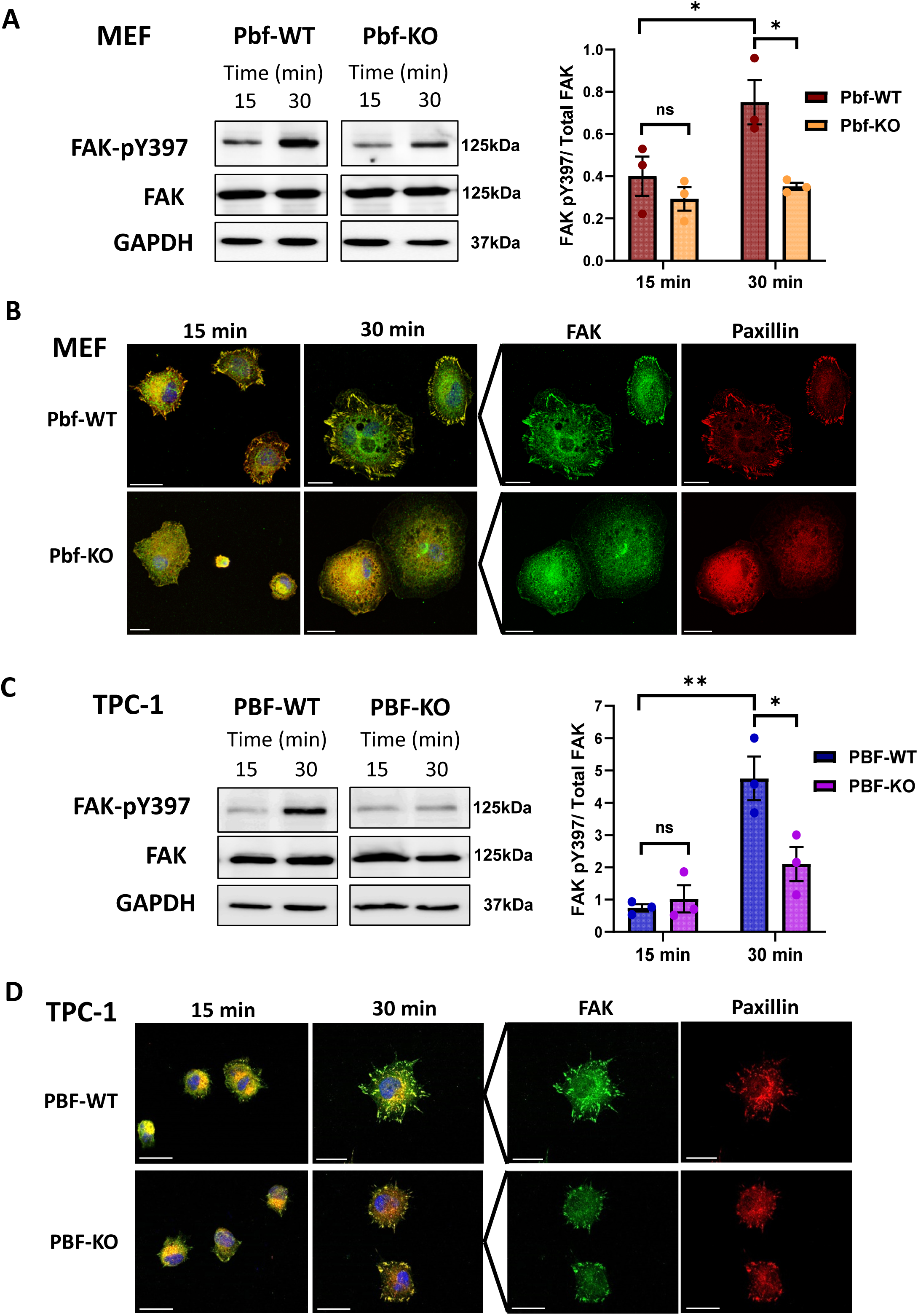
PBF loss delays FAK Y397 phosphorylation and early focal adhesion formation. **(A)** FAK Y397 phosphorylation in Pbf-WT and Pbf-KO MEFs 15 and 30 min after adhesion to fibronectin. FAK-pY397 levels were determined by Western blotting and normalised to total FAK. GAPDH was included as a loading control. **(B)** Representative immunofluorescence images of FAK (green) and paxillin (red) in Pbf-WT and Pbf-KO MEFs 15 and 30 min after adhesion to fibronectin. Merged images are shown on the left. **(C)** FAK Y397 phosphorylation in PBF-WT and PBF-KO TPC-1 cells 15 and 30 min after adhesion to fibronectin. FAK-pY397 levels were normalised to total FAK. **(D)** Representative immunofluorescence images of FAK and paxillin in PBF-WT and PBF-KO TPC-1 cells 15 and 30 min after adhesion to fibronectin. Merged images are shown on the left. N = 3–4 independent experiments. Data are presented as mean ± SEM. ns = not significant (*p* > 0.05), * = *p* < 0.05, ** = *p* < 0.01. Scale bars, 20 μm.

### PBF loss impairs front–rear polarity during directional migration

Front–rear polarity is critical for efficient directional migration in mammalian cells (1). To visualise the actin cytoskeleton in Pbf-KO MEFs during migration, we performed time-lapse microscopy following transfection with LifeAct-GFP. Pbf-WT MEFs exhibited a motile, spindle-like morphology with both filopodial and lamellipodial actin protrusions, particularly at the leading edge, prominent peripheral stress fibres and evidence of contractility and rear retraction (**Fig. 5A; Movie 1**). In contrast, Pbf-KO MEFs were more rounded, with broad lamellipodia present around the entire cell perimeter (**Fig. 5A; Movie 2**). Pbf-KO MEFs also displayed extensive cortical actin networks throughout the cell periphery with sustained retrograde flow (**Fig. 5A; Movie 2**).

**Figure 5.**
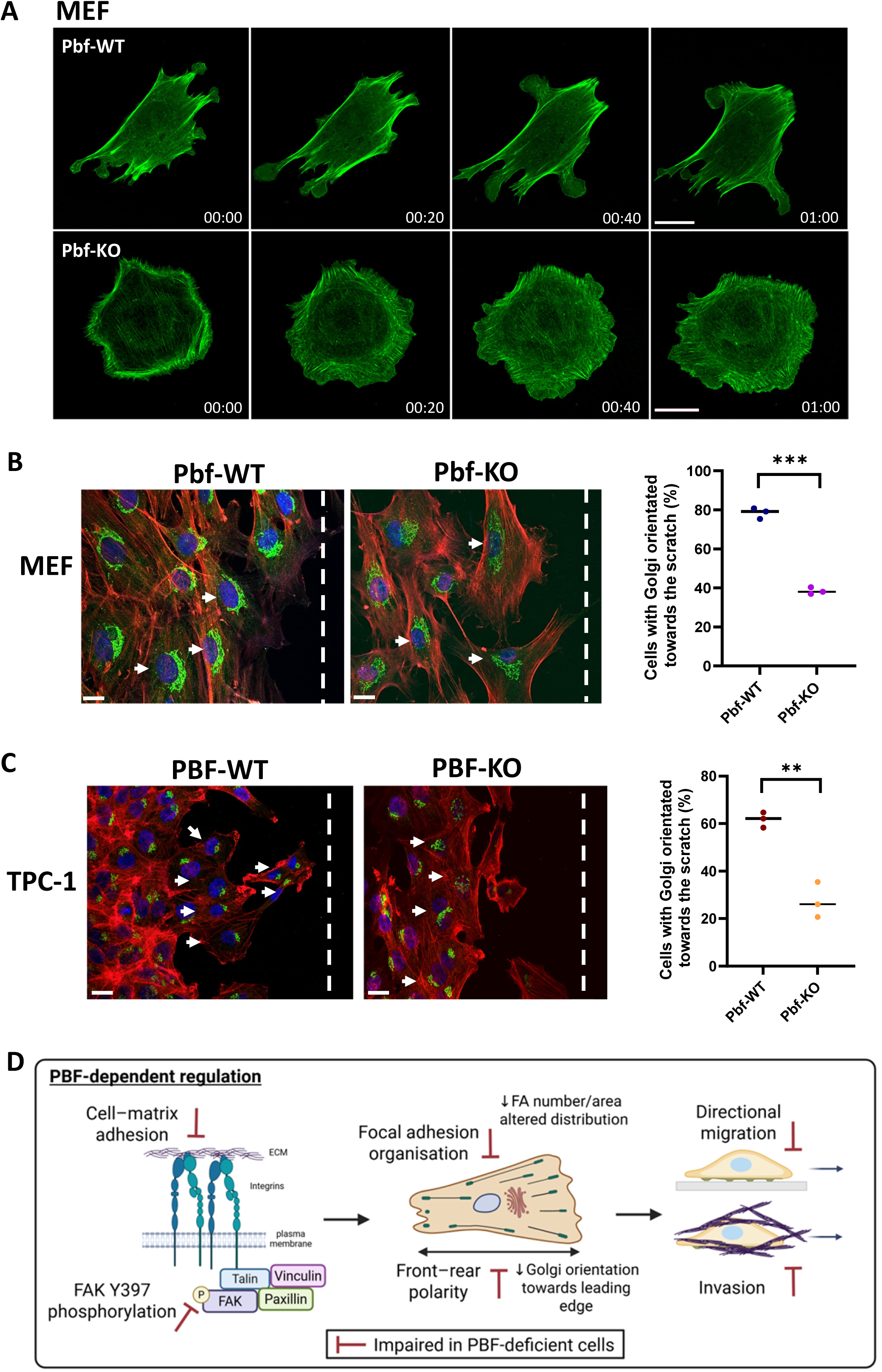
PBF loss impairs front–rear polarity during directional migration. **(A)** Time-lapse microscopy of LifeAct-GFP-expressing Pbf-WT and Pbf-KO MEFs migrating on fibronectin. Representative images are shown at 0, 20, 40 and 60 min. See also Movies 1 and 2. **(B)** Golgi orientation in Pbf-WT and Pbf-KO MEFs following scratch wounding. Representative images show GM130-labelled Golgi apparatus (green) and phalloidin-labelled F-actin (red). Quantification shows the percentage of cells with the Golgi orientated towards the scratch wound. **(C)** Golgi orientation in PBF-WT and PBF-KO TPC-1 cells following scratch wounding. Representative images show GM130-labelled Golgi apparatus (green) and phalloidin-labelled F-actin (red). Quantification shows the percentage of cells with the Golgi orientated towards the scratch wound. **(D)** Proposed model for PBF-dependent regulation of adhesion, polarity and directional motility. PBF supports a cell movement programme involving cell–matrix adhesion, FAK Y397 phosphorylation, focal adhesion organisation and front–rear polarity. In PBF-deficient cells, fibronectin adhesion and FAK Y397 phosphorylation are impaired, focal adhesions are reduced in number and area and show altered distribution, Golgi orientation is disrupted, and directional migration and invasion are reduced. Created in BioRender. Read, M. (2026) https://BioRender.com/1mrn3vj. For **(B)** and **(C)**, N = 3 independent experiments. Individual experimental values and means are shown. ** = *p* < 0.01, *** = *p* < 0.001. Scale bars, 20 μm.

These observations suggested defective front–rear polarity, which was assessed by determining Golgi orientation. During polarised migration, the Golgi apparatus is typically positioned between the nucleus and the leading edge (33). We therefore assessed Golgi orientation following scratch wound assays by immunofluorescent staining of the Golgi marker GM130. Significantly fewer Pbf-KO MEFs had the Golgi orientated towards the scratch wound, and GM130 staining was more diffuse and perinuclear compared with Pbf-WT MEFs (**Fig. 5B**). Similarly, PBF-KO TPC-1 cells demonstrated a significant reduction in Golgi orientation towards the scratch wound compared with PBF-WT TPC-1 cells (**Fig. 5C**).

These findings indicate that PBF-deficient cells are impaired in establishing front–rear polarity during directional migration. Taken together, our data identify PBF as a critical physiological regulator of cell movement, coordinating cell–matrix adhesion, focal adhesion signalling and polarity to support efficient migration and invasion (**Fig. 5D**).

## DISCUSSION

PBF has been characterised predominantly in cancer-associated settings, where pathological overexpression promotes tumourigenic phenotypes including proliferation, invasion and tumour formation (4, 6, 10, 11, 15, 17–26). However, whether these effects reflect an endogenous physiological role for PBF, or simply result from aberrant expression, has remained unclear. Here, we combine unbiased transcriptomic and phosphoproteomic profiling with genetic loss-of-function models to identify PBF as a regulator of coordinated cell movement. Altered PBF expression was associated with molecular networks linked to cell adhesion, ECM organisation, cytoskeletal regulation and cellular morphogenesis, while loss of endogenous PBF impaired adhesion, focal adhesion organisation, FAK phosphorylation, directional migration and invasion. These findings establish a physiological role for PBF in regulating adhesion-associated cell behaviour.

A key feature of this study is the convergence between transcriptomic and phosphoproteomic datasets. PBF overexpression altered expression of genes associated with cell adhesion, ECM organisation and cellular morphogenesis, suggesting that PBF influences the structural framework required for motility. In parallel, phosphoproteomic profiling identified changes in proteins involved in cytoskeletal organisation, cell polarity, adhesion and small GTPase signalling, indicating that altered PBF expression is associated with post-translational signalling networks that regulate dynamic cellular remodelling. The enrichment of motility-associated processes across both omics approaches suggested that cell movement may represent a major cellular process responsive to altered PBF expression, rather than a secondary or isolated observation. Importantly, these discovery analyses directed subsequent functional studies, which confirmed a requirement for endogenous PBF in migration and invasion.

The use of primary MEFs derived from a novel Pbf knockout mouse model allowed us to assess PBF function in a physiological cellular context. Pbf-deficient MEFs exhibited markedly impaired migration and invasion, while the intermediate phenotype observed in Pbf-HET MEFs suggests that PBF may regulate motility in a dose-dependent manner. Such dosage sensitivity is consistent with a role for PBF in coordinating multi-component cellular processes, where partial reduction may disrupt adhesion or signalling thresholds required for efficient movement. Targeted re-expression of PBF restored migration in Pbf-KO MEFs, supporting the specificity of the phenotype and arguing against nonspecific consequences of gene editing or cellular adaptation.

Our findings indicate that impaired motility in PBF-deficient cells is closely associated with defective adhesion. Both primary Pbf-KO MEFs and human PBF-KO TPC-1 cells showed reduced adhesion to fibronectin. These observations are consistent with the enrichment of adhesion-related GO terms in the omics datasets and suggest that PBF contributes to early cell–matrix interactions. The phenotypic convergence across primary and transformed models supports a conserved role for PBF in promoting efficient cell attachment.

Consistent with this, PBF loss profoundly altered focal adhesion organisation. In Pbf-KO MEFs, vinculin- and paxillin-positive adhesions were fewer, smaller and redistributed around the cell periphery. Similar alterations were observed in PBF-KO TPC-1 cells, indicating that the effect is not restricted to mouse fibroblasts. These changes occurred without major reductions in total vinculin, paxillin or FAK protein levels, suggesting that PBF does not primarily regulate abundance of these adhesion components. Instead, PBF appears to influence their spatial organisation and activation state. In support of this, FAK phosphorylation at Y397 was significantly reduced in both Pbf-KO MEFs and PBF-KO TPC-1 cells, whereas paxillin Y118 phosphorylation was not significantly altered. FAK Y397 autophosphorylation is a central event following integrin engagement and focal adhesion assembly (32), and its attenuation in PBF-deficient cells suggests that PBF is required for efficient early adhesion-associated FAK activation.

The early adhesion assays further support this interpretation. In Pbf-WT MEFs and PBF-WT TPC-1 cells, adhesion to fibronectin induced FAK Y397 phosphorylation and early peripheral adhesion formation. By contrast, PBF-deficient cells showed delayed or attenuated FAK activation and reduced early adhesion formation. These data position PBF upstream of, or parallel to, early adhesion-associated FAK activation and focal adhesion maturation. However, we do not yet establish whether PBF acts directly on FAK, on integrin activation or clustering, or through membrane trafficking pathways that regulate the delivery or organisation of adhesion-associated proteins. Further work will therefore be required to define the molecular step through which PBF regulates early adhesion signalling.

The focal adhesion defects observed in PBF-deficient cells may also be relevant to mechanosensing. Focal adhesions act as mechanosensitive hubs that translate cues from the ECM environment into cytoskeletal organisation, cell polarity and motility (34). In fibroblasts, matrix rigidity promotes polarisation via large, directionally organised adhesions, whereas cells on softer substrates remain radial, with small peripheral adhesions (34). Because focal adhesion alignment can precede cell elongation, the altered size and radial distribution of adhesions in PBF-deficient cells may reflect impaired matrix sensing and contribute to defective front–rear polarity.

In line with this, PBF deficiency disrupted cellular polarity. Live-cell imaging of LifeAct-GFP-expressing MEFs revealed striking differences in actin organisation: whereas Pbf-WT cells adopted an elongated, motile morphology with leading-edge protrusions and rear retraction, Pbf-KO cells were rounded and displayed broad circumferential lamellipodia with persistent cortical actin flow. This phenotype suggested a failure to establish or maintain stable front–rear polarity. Supporting this interpretation, Golgi orientation towards the scratch wound was significantly reduced in both Pbf-KO MEFs and PBF-KO TPC-1 cells. These observations suggest that PBF is required not only for adhesion formation, but also for the spatial coordination of cytoskeletal remodelling and polarity cues that enable directional migration. The enrichment of phosphoproteomic terms associated with small GTPase signalling is notable in this context, given the central role of Rho family GTPases in regulating protrusion, contractility and polarity (3). Although GTPase activity was not directly measured here, these findings provide a rationale for future studies examining whether PBF influences Rac1, RhoA or Cdc42 signalling during adhesion and migration.

A central question arising from this work is how PBF mechanistically links membrane-associated processes to adhesion and migration. Previous studies have shown that PBF can regulate the subcellular localisation of membrane proteins, including NIS and MCT8, promoting their redistribution from the plasma membrane to intracellular vesicular compartments (5, 27, 28). Our findings raise the possibility that endogenous PBF may similarly regulate proteins involved in cell–matrix adhesion or motility. For example, PBF could influence integrin trafficking, receptor recycling, adhesion complex assembly or spatial organisation of signalling molecules at the cell surface. Such a model would provide a mechanistic bridge between the established trafficking functions of PBF and the adhesion, FAK phosphorylation and polarity defects observed here. Future studies examining integrin localisation, activation and recycling will therefore be important to define how PBF regulates early adhesion signalling.

In summary, this study identifies PBF as a key physiological regulator of cell adhesion, polarity and directional motility. These findings extend the known functions of PBF beyond pathological overexpression and cancer-associated phenotypes, revealing an endogenous role in coordinating the adhesive and cytoskeletal processes that support cell movement. The viability of Pbf-KO mice suggests that PBF is not essential for survival under baseline conditions, but its importance may become more apparent in contexts requiring active cell migration, tissue remodelling or repair. More broadly, our data suggest that PBF links membrane-associated regulatory processes with the cellular machinery required for migration, providing a framework for future studies into its role in tissue organisation, repair, inflammatory remodelling and disease-associated invasion.

## MATERIALS AND METHODS

### Cell lines

Nthy-ori 3-1 thyroid follicular epithelial cells were obtained from the European Collection of Authenticated Cell Cultures (ECACC). TPC-1 papillary thyroid carcinoma cells were provided by Dr. Rebecca Schweppe (CU Cancer Center Tissue Culture Shared Resource, University of Colorado, Denver, USA). All cell lines were maintained in RPMI 1640 medium supplemented with 10% FBS, penicillin (10^5^ U/L), and streptomycin (100 mg/L) with regular mycoplasma testing and authentication. Nthy-ori 3-1 cells stably expressing untagged full-length PBF cDNA (4, 5) were generated through transfection of linearised plasmid (BglII-digested pCI-neo vector-only (VO) control and pCI-neo_PBF) using TransIT^®^-LT1 (Mirus Bio). Transfected cells were selected and maintained in 1 mg/mL G418.

CRISPR–Cas9 PBF knockout TPC-1 cells were generated as described elsewhere (13).

### Pbf-KO mouse model

C57BL/6NTac-Pttg1ip^em1(IMPC)H^/H mice (Pbf-KO) were obtained from the Mary Lyon Centre at MRC Harwell, which is the UK node of the European Mouse Mutant Archive (EMMA) (www.infrafrontier.eu; Repository number EM:11426). Pbf-KO mice were generated through CRISPR/Cas9-mediated deletion of *Pbf* exon 4 (ENSMUSE00001232242) (35, 36). All animals were housed under barrier conditions in ventilated cage racks in the Biomedical Services Unit (BMSU) at the University of Birmingham, which is maintained under artificial lighting between 07:00 and 19:00 (12 h), with controlled ambient temperature (22 °C ± 2 °C) and relative humidity (45%–65%). All animal experiments were performed in accordance with the Animals (Scientific Procedures) Act 1986 and UK Home Office regulations following approval by the local Animal Welfare and Ethical Review Body (AWERB).

### Pbf-KO mouse embryonic fibroblasts (MEFs)

Heterozygote timed matings were carried out to obtain embryos at embryonic day 13.5 and allow isolation of wild-type (WT), heterozygote (HET) and homozygote knockout (KO) mouse embryonic fibroblasts (MEFs). Where possible, within each experiment MEFs originated from the same litter.

Following removal of the brain, tail and internal organs, embryos were digested in 0.25% Trypsin/EDTA at 37 °C for 10 min and cells dissociated using an 18-gauge needle. Cells were grown on 0.2% (w/v) gelatin in DMEM High Glucose media containing 10% (v/v) FBS, 10^5^ U/L penicillin/100 μg/mL streptomycin, 2 mM L-Glutamine and 100 µM β-Mercaptoethanol and used in subsequent experiments before passage number 3.

To determine the genotype of each embryo, either tail or liver were digested in 10 mM Tris-HCl (pH 8.0), 25 mM EDTA, 50 mM NaCl, 0.5% SDS and 0.5 mg/mL Proteinase K at 55 °C overnight. Proteinase K was inactivated at 95 °C for 30 min and debris removed from the DNA solution by centrifugation at 1300 rpm for 1 min. The region flanking Pbf exon 4 was PCR amplified using the forward primer 5’-TGTAGACACTGGCTGAAAGG-3’ and reverse primer 5’-ACTCCATCATTACAGGCTGG-3’. Genotypes were analysed using agarose-gel electrophoresis.

MEFs were transfected in 6-well plates with HA-tagged PBF (PBF-HA) or pcDNA3.1+ vector-only (VO) control (4, 5) using Lipofectamine^TM^ 3000 (Invitrogen, L300015). Following the manufacturer’s instructions, 2 μg plasmid DNA was transfected with 5 μL P3000^TM^ Reagent and 5 μL Lipofectamine and cells were incubated for 24 h at 37 °C prior to protein extraction or wound healing assay.

### RNA-Seq analysis

RNA was extracted from Nthy-ori 3-1 cells stably expressing VO control and PBF using TRI Reagent™ (Sigma) and the RNeasy Micro Kit (QIAGEN), incorporating DNase I treatment. RNA sequencing (RNA-Seq) was performed by the Genomics Birmingham Genomics Service (University of Birmingham, UoB). RNA quantity and quality were assessed using the Qubit RNA HS Assay Kit (Invitrogen) and HS RNA ScreenTape assay (Agilent), respectively. Libraries were prepared using the QuantSeq 3’ mRNA-Seq Library Prep Kit FWD (Lexogen), and sequenced on a NextSeq 500 using a NextSeq 500/550 Mid Output Kit v2.5 (Illumina). Read quality was assessed using FastQC (37), adapters trimmed with Trim Galore! (v 0.5.0) (38) and reads mapped to the GRCh38 Human reference genome using STAR aligner (version 2.6.1b) (39). Gene-level counts were generated from binary alignment map (BAM) files using the Python library HTSeq (v 0.9.0) (40) and the featureCounts function in the R package Rsubread (v1.2.1) (41). Count normalisation and differential expression analysis were performed using DESeq2 (42), with p-values adjusted for multiple testing using the Benjamini-Hochberg FDR method.

### Phosphoproteomics analysis

Nthy-ori 3-1 cells stably expressing PBF were snap frozen in five T75 flasks, along with equivalent VO controls, and sent to Evotec AG (Munich) for phosphoproteomics analysis. Briefly, cells were lysed in urea-containing buffer and extracted proteins were proteolytically cleaved with endoproteinase Lys-C and trypsin. Peptides from each sample were isotopically encoded using isobaric tandem mass tags (TMT) and pooled into one sample. Combined peptides were fractionated by high-pH reversed phase chromatography into 12 fractions to decrease sample complexity and optimise phosphoproteome coverage. Each peptide fraction was enriched for phosphopeptides by immobilised metal affinity chromatography (IMAC) prior to liquid chromatography coupled to tandem mass spectrometry (LC-MS/MS) analysis. Mass spectrometric raw files were processed using the MaxQuant software package (43) for phosphopeptide/protein identification and TMT-based quantification. Significantly regulated phosphosites were determined by statistical testing based on the quantitative MaxQuant output files. To correct for multiple comparisons, p-values were adjusted using the Benjamini-Hochberg false discovery rate (FDR) correction procedure. Further details are provided in **Supplementary Methods**.

### Gene and phosphoprotein functional enrichment analysis

Transcriptomic and phosphoproteomic datasets were analysed separately for functional enrichment using ToppGene (44). Differentially expressed genes and proteins containing significantly altered phosphorylation sites were selected using thresholds of adjusted p < 0.1 and absolute fold change ≥ 1.2. For the phosphoproteomics analysis, quantitative data from 25,826 phosphorylation sites were included in statistical testing. The 5,879 proteins represented by these sites comprised the background set for enrichment analysis. For the transcriptomic analysis, 14,879 genes included in differential expression testing comprised the background set, with corresponding Ensembl Gene identifiers used for enrichment analysis. GO terms with Benjamini–Hochberg-adjusted p < 0.05 were considered significantly enriched. Enriched terms were consolidated into functional clusters according to gene or protein overlap, Jaccard similarity and hierarchical relationships within the Gene Ontology framework, thereby reducing redundancy and identifying coherent functional modules. For visualisation, the union count represents the total number of unique genes or proteins contained within each cluster. Statistical significance and enrichment values are shown for the most significant Gene Ontology term within each cluster, defined as the term with the lowest adjusted p value. Complete lists of significantly enriched terms are provided in the Supplementary Data.

### Antibodies and reagents

The following antibodies were used: rabbit polyclonal anti-PBF antibody (CSB-PA060105; Cusabio), rabbit polyclonal anti-PBF antibody (raised against the synthetic peptide RKKYGLFKEQNPYEKF [murine Pbf residues 159–174]; Covalab), mouse monoclonal anti-vinculin antibody (V9131; Sigma), rabbit polyclonal anti-paxillin antibody (PA5-34910; Invitrogen), mouse monoclonal anti-paxillin Y118 antibody (SC-365020; Santa Cruz), mouse monoclonal anti-FAK antibody (34Q36; Invitrogen), rabbit monoclonal anti-FAK-pY397 antibody (700255; Invitrogen), rabbit polyclonal anti-GM130 antibody (GTX130351; GeneTex), rabbit monoclonal anti-GAPDH antibody (D16H11; Cell Signaling Technology), and mouse monoclonal anti–β-actin antibody (clone AC-15; Sigma).

### Western blotting

Western blotting was performed as described previously (28). Proteins (30 µg per sample) were separated by SDS-PAGE. Membranes were probed with anti-PBF (Cusabio; 1:200), anti-vinculin (V9131; 1:1000), anti-paxillin (PA5-34910; 1:1000), anti-paxillin-pY118 (SC-365020; 1:1000), anti-FAK (34Q36; 1:500), anti-FAK-pY397 (700255; 1:500), anti-GAPDH (D16H11; 1:2000), and anti-β-actin (clone AC-15; 1:10,000) antibodies.

### Cell proliferation assay

To determine cell proliferation, the Cell Counting Kit-8 (CCK-8) assay was used in accordance with the manufacturer’s protocol (Sigma, 96992). Cells (4×10^3^ MEFs) were initially seeded into a 96-well plate with 3 technical replicates for each time point. At 0, 24 and 48 h cell media was discarded and 100 μL CCK8 solution (10 μL CCK8 + 90 μL complete media) was added to each well and incubated for 2 h at 37°C. Absorbance at 450 nm was measured using a plate reader (SpectraMax ABS Microplate Reader) and normalised against a complete medium only control. Subsequently, growth curves were generated.

### Wound healing assay

MEFs (8×10^5^) were seeded in a 6-well plate and cultured until they formed a confluent monolayer. A sterile pipette tip (200 μl) was used to create a wound in the cell monolayer and a DPBS wash removed any cell debris. Fresh culture medium was added to each well and a reference image of the scratch was taken immediately (0 h time point) using an EVOS XL Core Imaging System (ThermoFisher) with 10× magnification. Cells were maintained in standard growth conditions and further images were taken at 6, 12, and 24 h. The area of the wound was defined manually and measured within Fiji (45), and the percentage of wound recovery calculated by comparison to the initial (0 h) scratch area.

### Transwell migration and invasion assays

MEFs were seeded in a 6-well plate and grown until ∼80-90% confluent. Following serum starvation in medium containing 2% FBS for 4 h, the cells were trypsinised and reseeded in duplicate into the upper chamber of a 24-well plate containing Transparent PET membrane inserts with 8.0 μm pores (Falcon, 353097) for migration assays or BioCoat™ Growth Factor Reduced Matrigel Invasion Chambers (Corning, 354483) for invasion assays. 2×10^4^ MEFs were seeded into the upper chamber of each well in media containing 2% FBS while media containing 10% FBS was added to the lower chamber to act as chemoattractant for the migrating cells. After incubation at 37 °C for 24 h, the migrated cells were fixed in 95% ethanol for 5 min and stained with Mayer’s haematoxylin (MHS16) for 10 min and Eosin (HT110116) for 5 min. Cells were dehydrated in 70% and 90% ethanol and air dried before examination under a Leica DM2000 microscope at 10× magnification. Five fields of view were imaged per insert and the total number of migrated cells determined using manual counting in Fiji (45).

### Cell adhesion assays

Pbf-WT, Pbf-HET and Pbf-KO MEFs (8×10^4^) or parental and PBF-KO TPC-1 cells (6×10^4^) were seeded into 6-well plates containing coverslips coated with fibronectin (10 µg/mL; Sigma, F0895). After a 30 min or 1 h incubation at 37 °C, non-adherent cells were removed and adherent cells were fixed in 4% paraformaldehyde. Following blocking with 0.1 M glycine, cells were permeabilised with 0.1% Triton X-100 and then incubated with Alexa Fluor 594-phalloidin (Invitrogen, A-12381) diluted 1:40 in 0.1% Triton X-100 for 1 h at room temperature. Subsequently, cells were washed three times with 0.1% Triton X-100 and mounted using ProLong Gold Antifade Mountant with DAPI (Invitrogen, P36931). Phalloidin-stained cells were imaged using a Zeiss Axio Observer widefield system with Apotome 2.0 technology. For each time point, 12-15 images captured from different areas of the slide were analysed in each of 3-4 biological repeats. Cell segmentation was achieved using a modified version of a platelet spreading analysis previously described (46) with a customised Cellpose 2.0 model (47, 48). This was achieved by fine-tuning the “cyto2” model on a subset of the data before running the customised model on the full dataset. A Jupyter notebook (49) was then used to quantify adherent cell numbers. Both notebooks are available at https://github.com/JeremyPike/cell-classification.

### Immunofluorescence (IF) staining

Pbf-WT and Pbf-KO MEFs (5×10^4^) or parental and PBF-KO TPC-1 cells (4×10^4^) were seeded onto coverslips coated with fibronectin (10 µg/mL). After 24 h, cells were fixed in 4% paraformaldehyde and permeabilised with 0.1% saponin before incubation with primary antibodies for 1 h (anti-vinculin [V9131], 1:100; anti-paxillin [PA5-34910], 1:100), followed by incubation with secondary antibodies and probes (Alexa Fluor 594-phalloidin, 1:40; Alexa Fluor 488-conjugated goat anti-mouse [A-11001], Alexa Fluor-488-conjugated goat anti-rabbit [A-11008], and Alexa Fluor 555-conjugated goat anti-rabbit [A-21428] (Invitrogen), all at 1:250). Finally, coverslips were mounted onto slides using ProLong Gold Antifade Mountant with DAPI and images captured on an LSM880 Airyscan confocal microscope using a 40× or 100× objective.

### Focal adhesion quantification

Pbf-WT and Pbf-KO MEFs were stained with vinculin for quantification of focal adhesions. Images were captured on an LSM880 Airyscan confocal microscope using a 40× objective. A custom Fiji macro (45) was used to batch process the images. For each image, noise was reduced using a Gaussian filter with a standard deviation of 2 pixels. Background signal was subtracted using a rolling ball approach with a ball radius of 15 pixels. Subsequently, the contrast of the images was enhanced and an automated Li threshold applied to produce a binary segmentation image. Focal adhesions were defined as connected components in the binary image larger than 150 pixels. Cell-level measurements of focal adhesion number, mean area and mean vinculin fluorescence intensity were calculated for a total of 102 Pbf-WT and 108 Pbf-KO MEFs across three independent experiments. These measurements were analysed using generalised linear mixed models to account for clustering of cells within experimental repeats.

### Early adhesion assays

Pbf-WT, Pbf-HET and Pbf-KO MEFs, or parental and PBF-KO TPC-1 cells (1×10^5^) were seeded into fibronectin-coated (10 µg/mL) 6-well plates with or without coverslips for IF and protein analysis, respectively. Following incubation at 37 °C for 15 or 30 min, non-adherent cells were removed, and adherent cells were analysed by IF following staining with anti-paxillin (PA5-34910; 1:100) and anti-FAK (34Q36; 1:50) antibodies, or by Western blotting to assess FAK phosphorylation at Y397.

### Time-lapse microscopy

Pbf-WT, Pbf-HET and Pbf-KO MEFs (5×10^4^) were seeded into 6-well plates coated with fibronectin (10 µg/mL) and incubated for 24 h. Cells were then transfected with pLifeact-TagGFP2 (kind gift from Dr Chris Weston, UoB). After an additional 24 h, the transfected cells were trypsinised and re-seeded into glass-bottom imaging dishes (VWR confocal dishes, Cat no: 734-2904) coated with fibronectin (10 µg/mL). Live-cell imaging was performed using a Zeiss LSM880 Airyscan confocal microscope equipped with a 40× objective, with time-lapse images acquired every 1 min for up to 1 h under physiological conditions (37 °C, 5% CO₂).

### Golgi orientation measurement

Cells were seeded in 6-well plates with coverslips coated with fibronectin (10 µg/mL) and a scratch was induced after 24 h as described for the wound healing assays. Following a 16 h incubation, cells were fixed and IF stained with anti-GM130 antibody (GTX130351; 1:100) and Alexa Fluor 594-phalloidin (1:40). Golgi orientation was measured following established methods whereby lines are placed over cell nuclei at a 120° angle to delineate the front 1/3 and rear 2/3 of the cell area with respect to the direction of migration (50, 51). Cells were defined as having Golgi orientated towards the scratch wound if the majority (at least 50%) of the Golgi apparatus was localised within the 120° sector facing the wound edge. N=73-157 cells per condition were analysed across three independent experiments.

### Statistical analysis

Data were analysed using IBM SPSS Statistics (version 29). Normality of data was assessed using the Shapiro-Wilk test. Student’s *t*-test and Mann-Whitney U test were used for comparison between two groups of parametric and nonparametric data, respectively. One-way ANOVA with Tukey’s post-hoc test and a Kruskal-Wallis test with Dunn’s post-hoc test were used for comparisons of multiple groups of parametric and nonparametric data, respectively. Cell-level focal adhesion measurements were analysed using generalised linear mixed models with genotype as a fixed effect and experimental repeat as a random effect. Significance was taken as *p* < 0.05.

## ACKNOWLEDGEMENTS

Mice used in this study were obtained from the Mary Lyon Centre at MRC Harwell (MLC) and the following award is acknowledged: MC_UP_2201/2. The MLC is also a member of the International Mouse Phenotyping Consortium (IMPC) and has received funding from the National Institutes for Health [5UM1HG006348-08] for generating and/or phenotyping the C57BL/6NTac-Pttg1ip^em1(IMPC)H^/H mice.

We acknowledge the support of University of Birmingham Facilities: the BMSU staff for expert animal husbandry, the Microscopy Facility (RRID:SCR_027108) for providing access to equipment and technical expertise, and the Genomics Birmingham Facility for supporting the RNA-Seq experiments.

We also thank the technical and scientific staff at Evotec AG (Munich) for their support with quantitative phosphoproteome analysis, the CU Cancer Center Tissue Culture Shared Resource, which receives direct funding support from the National Cancer Institute through the Cancer Center Support Grant (P30CA046934), for access to authenticated cell lines and Chris Weston (University of Birmingham) for providing the pLifeact-TagGFP2 plasmid.

## DECLARATION OF GENERATIVE AI IN THE WRITING PROCESS

During preparation of this manuscript, Microsoft Copilot (https://m365.cloud.microsoft/chat) was used to assist with improving the clarity, grammar, and readability of the text. The authors critically reviewed and edited all AI-assisted text and take full responsibility for the accuracy, integrity and final content of the manuscript.

