## Supplementary Information for "PBF/PTTG1IP coordinates focal adhesion formation, polarity and cell motility"

**Supplementary Figure 1**

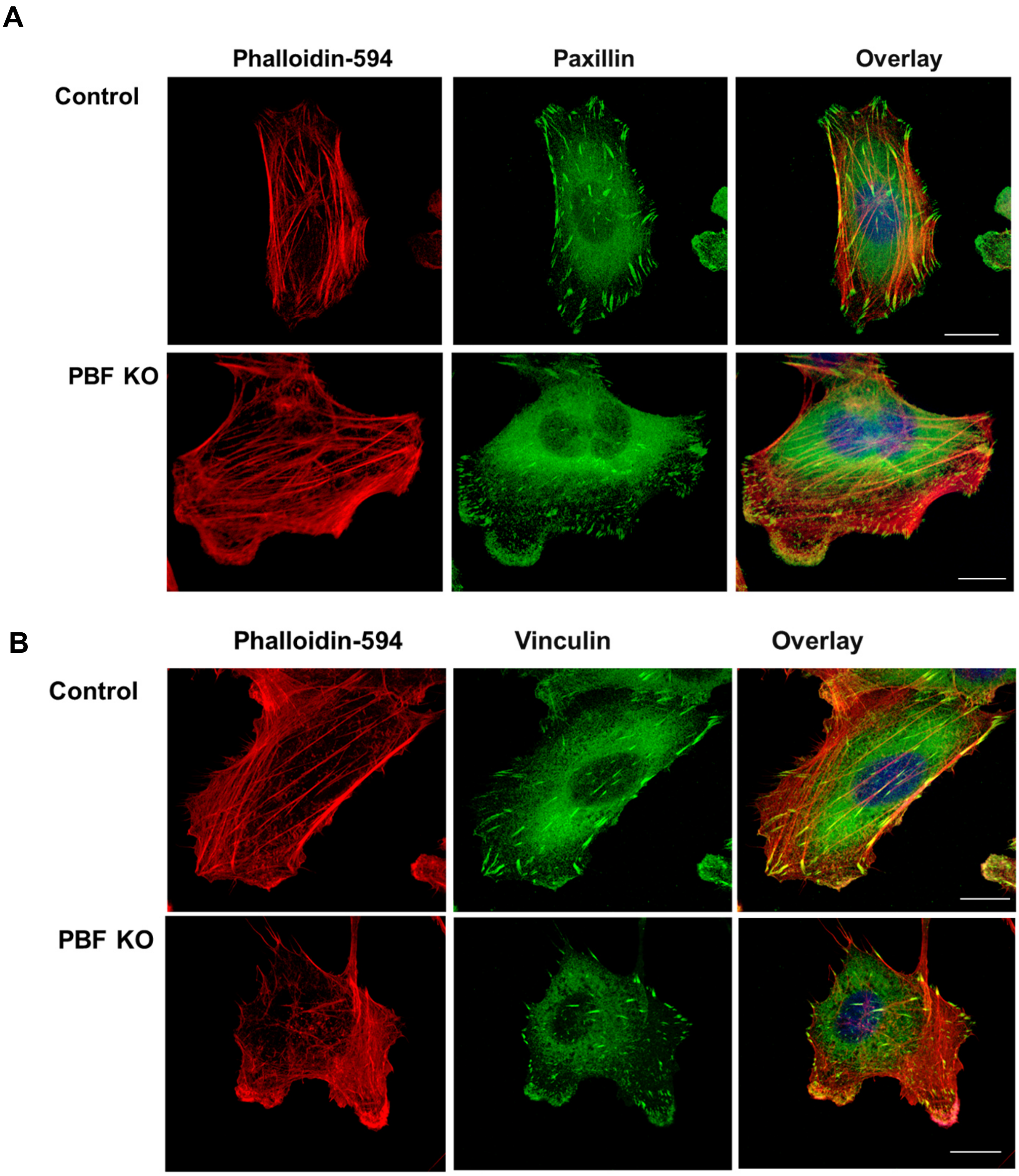


**Supplementary Figure 2. Focal adhesion analysis in TPC-1 PBF knockout (KO) cells.** Immunofluorescent staining of PBF-WT and PBF-KO TPC-1 cells with focal adhesion markers paxillin (**A**) and vinculin (**B**) shown in green and phalloidin-594 in red. Representative images shown (N=3). Bars = 20μm.

### Supplementary Methods

#### Phosphoproteomic analysis

##### MS sample preparation

Cells were lysed in 300 µL ice-cold urea-containing buffer (8 M urea, 50 mM Tris-HCl pH 8.2, 75 mM NaCl, 5 mM EDTA, 5 mM EGTA, 10 mM sodium pyrophosphate, 10 mM glycerol phosphate, 10 mM sodium fluoride, 2.5 mM sodium orthovanadate, protease inhibitor cocktail Complete Mini (Roche), phosphatase inhibitor cocktails 2 and 3 (Sigma)), and sonicated three times for 1 minute on ice. Protein concentrations were determined (Bradford assay, Bio-Rad) and equal protein amounts (0.2 mg) of each of the 10 samples (5x VO, 5x PBF) were reduced by dithiothreitol and alkylated by iodoacetamide, followed by digestion with LysC and Trypsin (Promega). Proteolytic peptides were then desalted using reversed-phase 100 mg $C_{18}$ SepPak cartridges (Waters) followed by chemical TMT 10-plex labelling (ThermoFisher) according to the manufacturer's instructions, with labelling efficiencies of >95% confirmed. Samples were pooled and a total of 2 mg peptide was fractionated by high pH reversed phase chromatography. Peptides were reconstituted in 20 mM ammonium formate (pH 10, buffer A), loaded onto a XBridge $C_{18}$, 200 × 4.6 mm column (Waters) operated with the Äkta Explorer system (GE Healthcare) and separated by applying a segmented gradient increasing the acetonitrile concentration from 7% to 30% buffer B (buffer A supplemented with 80% acetonitrile) over 15 min followed by a 5 min gradient to 55%. Fractions were combined in a non-linear way to generate 12 samples with equal peptide amounts that were subsequently frozen in liquid nitrogen, lyophilised, reconstituted in 0.1% TFA and desalted using 100 mg $C_{18}$ Sep-Pak columns (Waters).

Phosphopeptides were enriched using immobilised metal affinity chromatography (IMAC). Peptides of each fraction were reconstituted in IMAC loading buffer (80% acetonitrile containing 0.1% TFA) at a final concentration of approximately 5 mg/ml. To generate the IMAC-resin, Ni ions of Ni-NTA Superflow Agarose Beads (Qiagen) were removed by incubation with 100 mM EDTA and subsequently replaced by $Fe^{3+}$ ions. The resin was then washed with $H_2O$ and reconstituted in a 1:1:1 mix of acetonitrile, methanol and 0.01% acetic acid. To each sample, 10 µL of equilibrated IMAC resin was added and incubated for 30 min at 25°C and 1,400 rpm in a Thermomixer (Eppendorf). This slurry was then loaded onto in-house build $C_{18}$ STAGE tip columns. Subsequently, beads were washed with IMAC loading buffer and phosphopeptides were eluted onto the $C_{18}$ material by washing twice with a 500 mM $K_2HPO_4$ solution. The $C_{18}$-bound phosphopeptides were washed with 0.1% formic acid, eluted with 50% ACN, 0.1% FA, concentrated in a Vacufuge$^{TM}$ (Eppendorf) and reconstituted in 0.1% FA for MS analysis.

##### Mass spectrometric analysis

LC-MS/MS analyses were performed on a Q Exactive HF mass spectrometer (Thermo Fisher Scientific), equipped with an Easy n-LC 1000 UPLC system (Thermo Fisher). Samples were loaded with an auto sampler onto a 40 cm self-made fused silica emitter packed with reversed phase material (Reprusil-Pur $C_{18}$-AQ, 1.9 µm, Dr. Maisch GmbH) at a maximum pressure of 950 bar. Bound peptides were eluted in 120 min run time and sprayed directly into the mass spectrometer using a nanoelectrospray ion source (Thermo Fisher).

The mass spectrometer was operated in a data-dependent acquisition mode to automatically switch between full scans (resolution R=60.000 at m/z 400) and the acquisition of HCD fragmentation spectra (R=30.000 at m/z 400) of the ten (max. IT=100 msec, AGC target=2e5) most abundant peptide ions using an isolation window of 1.4 m/z. Peptides selected once for fragmentation were dynamically excluded for 30 sec for further fragmentation.

**Data processing and statistical analysis**

The obtained raw files were processed with the MaxQuant software suite for peptide identification and quantification using the human Uniprot database (version 09 2017). Carbamidomethylation of cysteine was set as a fixed modification and oxidation of methionine, N-terminal acetylation and the phosphorylation of serine, threonine and tyrosine residues were set as variable modifications. All peptides were required to have a minimum peptide length of seven amino acids and a maximum of two missed cleavages and three modified amino acids were allowed. The false discovery rate (FDR) for peptide identifications was limited to 1% based on a target-decoy approach. For quantification, the intensities of TMT reporter ions were used.

The MaxQuant Phospho(STY)Sites.txt files were used to analyse the data. Features flagged as decoys or potential contaminant were excluded. The MaxQuant-corrected TMT reporter intensities were $log_{10}$-transformed and scaled so that the median intensity was equivalent across samples.

Differentially regulated phosphosites were identified by one-way ANOVA with experimental group as the factor. The obtained p-values were corrected for multiple testing using the Benjamini–Hochberg procedure. Only phosphosites quantified in at least two-thirds of replicates within each experimental group were included in statistical testing.
